# The affinity of the S9.6 antibody for double-stranded RNAs impacts the mapping of R-loops in fission yeast

**DOI:** 10.1101/217083

**Authors:** Stella R. Hartono, Amélie Malapert, Pénélope Legros, Pascal Bernard, Frédéric Chédin, Vincent Vanoosthuyse

**Author notes:** equal contribution (Stella R. Hartono, Amélie Malapert).

## Abstract

R-loops, which result from the formation of stable DNA:RNA hybrids, can both threaten genome integrity and act as physiological regulators of gene expression and chromatin patterning. To characterize R-loops in fission yeast, we used the S9.6 antibody-based DRIPc-seq method to sequence the RNA strand of R-loops and obtain strand-specific R-loop maps at near nucleotide resolution. Surprisingly, preliminary DRIPc-seq experiments identified mostly RNase H-resistant but exosome-sensitive RNAs that mapped to both DNA strands and resembled RNA:RNA hybrids (dsRNAs), suggesting that dsRNAs form widely in fission yeast. We confirmed *in vitro* that S9.6 can immuno-precipitate dsRNAs and provide evidence that dsRNAs can interfere with its binding to R-loops. dsRNA elimination by RNase III treatment prior to DRIPc-seq allowed the genome-wide and strand-specific identification of genuine R-loops that responded *in vivo* to RNase H levels and displayed classical features associated with R-loop formation. We also found that most transcripts whose levels were altered by *in vivo* manipulation of RNase H levels did not form detectable R-loops, suggesting that prolonged manipulation of R-loop levels could indirectly alter the transcriptome. We discuss the implications of our work in the design of experimental strategies to probe R-loop functions.

## INTRODUCTION

R-loops are three-stranded nucleic acid structures composed of a stable DNA:RNA hybrid and a displaced single-stranded DNA (Santos-Pereira and Aguilera 2015). R-loops have been found from bacteria to higher eukaryotes and usually form upon re-annealing of the nascent transcript to its DNA template in the wake of the RNA polymerase. R-loop formation is antagonized by a number of enzymes, including RNase H enzymes or DNA&RNA helicases such as the highly conserved Senataxin (reviewed in (Skourti-Stathaki and Proudfoot 2014)). In addition, nucleosome turnover (Herrera-Moyano et al. 2014; Taneja et al. 2017), chromatin structure and topology or the efficient packaging of nascent RNA into ribonucleotide particles (mRNPs) were also proposed to limit R-loop formation because they restrict the possibility of hybridization between the nascent RNA and its DNA template (reviewed in (Santos-Pereira and Aguilera 2015)).

R-loops are widely mapped genome-wide by exploiting the strong affinity of the S9.6 antibody for DNA:RNA hybrids (Chedin 2016). Recent maps have shown that R-loops can occupy between 5% and 10% of the genome in yeast, vertebrates and *Arabidopsis thaliana* (Sanz et al. 2016; Wahba et al. 2016; Xu et al. 2017). R-loop formation has been associated with replication stress and genome instability and it was recently proposed that defective control of R-loop formation could contribute to a number of human pathologies (reviewed in (Skourti-Stathaki and Proudfoot 2014; Santos-Pereira and Aguilera 2015; Richard and Manley 2016)). However, a mounting body of evidence also suggests that R-loops play physiological roles in transcriptional control and chromatin patterning (reviewed in (Skourti-Stathaki and Proudfoot 2014; Santos-Pereira and Aguilera 2015; Chedin 2016)). What distinguishes toxic and beneficial R-loops is still unclear. To answer this question, accurate maps of R-loops must be obtained and assays must be developed to sort R-loops into functional categories.

Whether the S9.6 antibody is the best tool to map R-loops has recently been debated. In-depth characterization of the S9.6 antibody *in vitro* has shown that it recognizes preferentially DNA:RNA hybrids but that it can also recognize AT-rich RNA:RNA hybrids (dsRNAs), albeit with a five-fold lower affinity (Phillips et al. 2013). In addition, it was argued that the affinity of S9.6 for R-loops is influenced by R-loop sequences (Konig et al. 2017). The direct consequences of these observations on the accuracy of R-loop mapping by DRIP-like methods have not yet been rigorously assessed. Preliminary RNase A treatment have been proposed to increase the specificity of R-loop mapping using DRIP approaches (Zhang et al. 2015) but this was later disputed (Halasz et al. 2017). However, whether or not RNase A treatment does modulate the efficacy of DRIP, it does not help to evaluate whether dsRNAs are able to interfere with the quantitative recovery of R-loops using S9.6, as RNase A will degrade single stranded RNA but not dsRNA. To date, it is still not clear whether S9.6 can immuno-precipitate dsRNAs *in vivo* or whether dsRNA can interfere with the interaction between S9.6 and DNA:RNA hybrids.

Here we sought to identify R-loops in the fission yeast *Schizosaccharomyces pombe* using the recently described DRIPc-seq approach, where RNA:DNA hybrids are immuno-precipitated with the S9.6 antibody and their RNA strands are sequenced in a strand-specific manner to map R-loops at near base-pair resolution (Sanz et al. 2016). This strategy is well suited to mapping R-loops in compact genomes such as fission yeast where genes frequently overlap. Surprisingly, we found that the presence of exosome-sensitive dsRNAs in fission yeast interfered with the mapping of R-loops using this method. Robust R-loop identification by DRIPc-seq required dsRNA elimination using RNase III treatment. We show that these newly mapped R-loops respond to RNase H levels and exhibit the expected characteristics of R-loop forming genes. Surprisingly, our results did not confirm that R-loops form over centromeric heterochromatin repeats, as suggested previously (Nakama et al. 2012; Castel et al. 2014). Our data indicate instead that S9.6 was able to immuno-precipitate RNase III-sensitive dsRNAs derived from centromeric hetrochromatic repeats. Taken together, our results demonstrate that the affinity of the S9.6 antibody for dsRNAs has the potential to interfere with the accurate and quantitative identification of R-loops by DRIP-like approaches. In addition, we provide evidence that long-term manipulation of RNase H levels causes a consistent transcriptional response affecting mostly non-R-loop forming genes.

## RESULTS

### The S9.6 antibody immuno-precipitates double-stranded RNAs (dsRNAs) *in vitro* and *in vivo*

To map R-loops in fission yeast, we used the recently described DRIPc-seq method, where RNA:DNA hybrids are immuno-precipitated with the S9.6 antibody and their associated RNA strands are sequenced in a strand-specific manner to map R-loops at near base-pair resolution (Sanz et al. 2016) (Figure S1). Unexpectedly, our first attempts at DRIPc-seq failed to deliver strand-specific signals. Instead, the vast majority of RNAs sequenced mapped to both DNA strands over genes in a nearly symmetrical pattern. This pattern was clearly distinct from the one obtained when sequencing total RNA, suggesting that DRIPc-seq enriched for a specific class of low-abundance RNAs (Figure 1A). Such DRIPc-seq RNAs were largely resistant to exogenous RNase H treatment and to *in vivo* expression of exogenous RNase H (*Escherichia coli* RnhA), strongly suggesting that they did not correspond to R-loops (Figure 1A and data not shown). For about half the genes, those RNase H-resistant DRIPc-seq RNAs were particularly enriched at gene ends (Figure 1A and see gene clusters #1-3, Figure S2), consistent with the idea that they only represent a specific subset of total RNAs. Interestingly, RNase H-resistant DRIPc-seq RNAs were both longer and more abundant when the activity of the RNA exosome was impaired by the deletion of Rrp6 (*rrp6*Δ) or the *dis3-54* mutation (Figure 1BCD and Figure S3). Because such RNase H-resistant and exosome-sensitive RNAs mapped to both DNA strands, we speculated that they were double-stranded RNAs (dsRNAs). Dot blot analysis with the dsRNA-specific J2 antibody confirmed that both exosome mutants accumulate dsRNAs (Figure 1E). Taken together, these observations suggested that our first attempts at DRIPc-seq in fission yeast identified mostly RNase H-resistant and exosome-sensitive dsRNAs but not R-loops.

**Figure 1.**
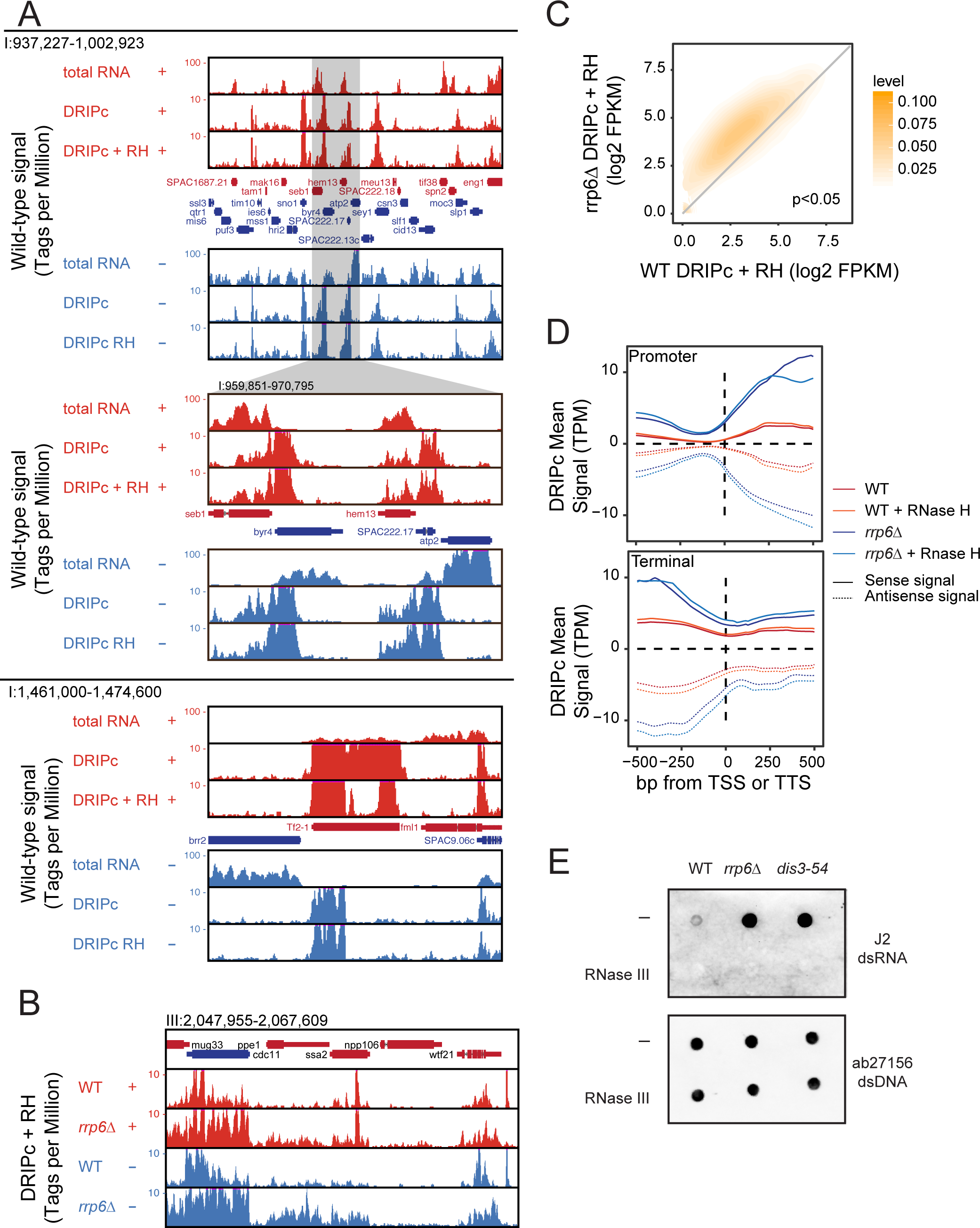
DRIPc-seq identifies RNase H-resistant but exosome-sensitive transcripts in fission yeast. **(A)** Representative genomic regions showing RNase H-resistant DRIPc-seq signals in wild-type cells. Signals are broken down by strand (red = + strand; blue = − strand) and represent total RNA, DRIPc-seq, and RNase H-pre-treated DRIPc-seq in each sub-panel. Annotated genes are shown. Note that RNase H-resistant DRIPc-seq RNAs are often enriched at regions of gene convergence and that DRIPc-seq RNAs at Tf2 transposons are partially sensitive to exogenous RNase H pre-treatment. **(B)** Representative genomic region showing RNase H-resistant DRIPc-seq signals in rrp6+ and *rrp6*Δ cells. **(C)** X-Y plot of RNase H-resistant DRIPc-seq signals at each locus in *rrp6*Δ (y-axis) vs. rrp6+ (x-axis) cells. **(D)** Metaplot of DRIPc-seq signals in rrp6+ and *rrp6*Δ cells at promoter and terminator regions **(E)** Nucleic acids extracted from wild-type or exosome-deficient *rrp6*Δ and *dis3-54* cells were pre-treated or not with RNase III and the amount of dsRNA or dsDNA was analyzed on separate dot blots.

It was reported previously that the S9.6 antibody can recognize AT-rich dsRNAs *in vitro*, albeit with a lower affinity than DNA:RNA hybrids (Phillips et al. 2013). Here we show that the S9.6 antibody is able to recognize and immuno-precipitate long GC-rich dsRNAs produced *in vitro* from the mouse *Airn* gene (Ginno et al. 2012) (Figure 2AB). The R-loop forming *Airn* gene encoded on a plasmid was transcribed *in vitro* either in the sense orientation (using T3 RNA polymerase) or in both sense and antisense orientation (using both T3 and T7 RNA polymerases). As described previously (Ginno et al. 2012), transcription in the sense orientation was associated with an R-loop-dependent topological shift of the plasmid on agarose gels, that was sensitive to RNase H but resistant to RNase III treatment (Figure 2Aa, lanes 2-5). Sense transcription also produced strong RNase H-sensitive and RNase III-resistant signals on S9.6 dot blots (Figure 2Ab, dots 2-5). Transcription in both orientations was associated with the production of RNase H- and RNase A-resistant but RNase III-sensitive double-stranded RNAs (dsRNAs) that ran on agarose gels as a smear (Figure 2Aa, lanes 6-9). Strikingly, the associated S9.6 dot blot signals were partially resistant to RNase H treatment (dot #7, Figure 2Ab) and only disappeared fully when treated with both RNase H and RNase III (dot #9, Figure 2Ab). This strongly suggested that S9.6 was able to recognize GC-rich dsRNAs produced from *Airn* as well as R-loops. This was confirmed by spotting only dsRNAs on S9.6 dot blots (dot #10, Figure 2Ab). To build on these observations, we tested whether S9.6 could immuno-precipitate dsRNAs produced from *Airn*. dsRNAs were incubated either alone or in combination with T3-transcribed *Airn* as a source of R-loops and immuno-precipitated with S9.6. We analysed the input (IN), the unbound fraction (FT), the last wash (W4) and the bound fraction (IP) using agarose gel (Figure 2Ba), dot blots (Figure 2Bb) and qPCR (Figure 2Bc) (see Methods). dsRNAs alone were efficiently captured by S9.6, whether or not R-loops were present. In the absence of dsRNAs, the vast majority of R-loops were captured by the S9.6 antibody since no R-loop was found in the unbound fraction (Figure 2Bb). Strikingly however, when dsRNAs were incubated with R-loops, some R-loops did not bind to S9.6 and remained in the unbound fraction (Figure 2Bb&2Bc). As a result, fewer R-loops were found in the bound fraction (Figure 2Bc). This experiment showed conclusively that S9.6 could immuno-precipitate dsRNAs and that dsRNAs were able to interfere with the association of S9.6 with R-loops, at least *in vitro* (Figure 2B).

**Figure 2:**
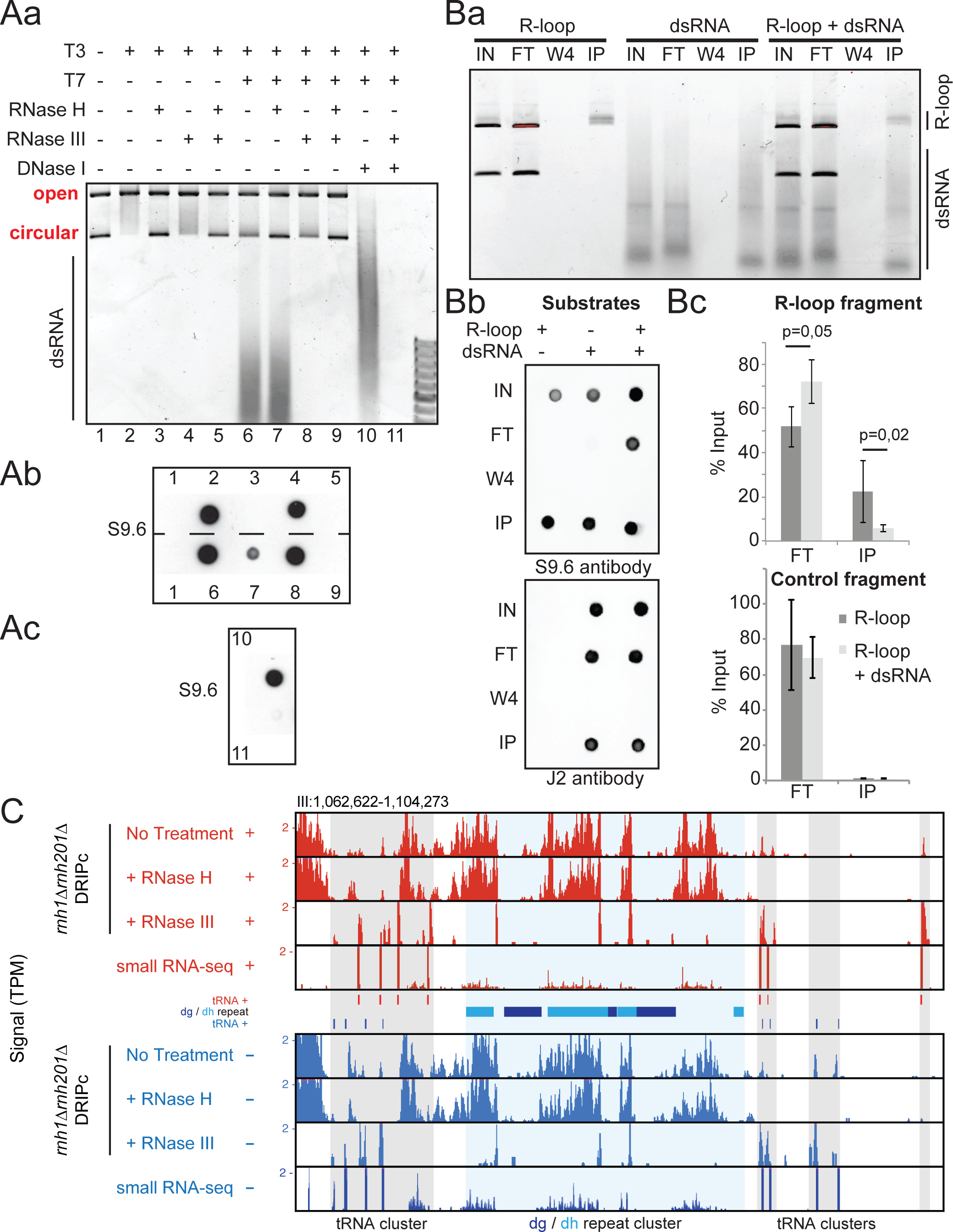
The S9.6 antibody recognizes and immune-precipitates double-stranded RNAs both *in vitro* and *in vivo* in fission yeast. **(A)** A plasmid carrying the R-loop forming *AIRN* mouse gene (Ginno et al. 2012) was transcribed *in vitro* in the sense (T3) or antisense (T7) orientations as indicated. Soluble RNAs and proteins were then digested with RNase A and Proteinase K, respectively. After concomitant sense and antisense transcription, the reaction was split in two. In one half of the reaction, the plasmid was digested using DNase I treatment and only the dsRNA were kept. **(Aa)** The reactions were run on an agarose gel. R-loops formed upon sense transcription (T3) induced a topological shift in the plasmid as reported previously (Ginno et al. 2012). When both T3 and T7 polymerases were used at the same time, RNase III-sensitive dsRNAs appeared in the gel. **(Ab)** Dot blot analysis of the above reactions using the S9.6 antibody. Upon sense and antisense transcription, the RNase H treatment is not sufficient to remove the signal. The signal fully disappears only after treatment with both RNase H and RNase III, suggesting that the S9.6 can recognize dsRNA *in vitro.* This was confirmed using the purified dsRNA **(Ac)**. **(B)** R-loops and dsRNA were produced from the pFC53 plasmid. After transcription, the R-loop containing plasmids were cut with ApaLI to produce 2 bands, the longest of which contains R-loops. The smallest band acts as negative control because it does not carry R-loops. The cut plasmids and dsRNAs were then incubated alone or in combination with 4 μg of purified S9.6 antibody. The input (IN), corresponding to half of what was incubated with the antibody, the flow-through fraction (FT) corresponding to the material that did not bind to the antibody, the last wash (W4) and the immuno-precipitated fraction (IP) were then purified using phenol/chloroform extraction and ethanol precipitation and either run on an agarose gel **(Ba)**, or analysed by dot blot with the indicated antibodies **(Bb)**. **(Bc)** The long fragment where R-loops can form (R-loop fragment) and the short fragment where R-loops do not form (control fragment) were also quantified by qPCR (n = 4 independent experiments, p-values with Wilcoxon - Mann Whitney). The S9.6 antibody can immuno-precipitate GC-rich dsRNAs and when dsRNAs are mixed with R-loops, the recovery of R-loops by the S9.6 antibody is reduced (see text). **(C)** Representative centromeric region of chromosome III showing RNase H-resistant but RNase III-sensitive DRIPc-seq signals at the *dg* and *dh* centromeric repeats. Note that most DRIPc-seq signals over tRNA genes are RNase H-sensitive, confirming that genuine R-loops form at tRNA genes (Legros et al. 2014).

To test whether S9.6 can also recognize dsRNA produced *in vivo*, we studied the centromeric *dg* and *dh* repeats. These repeats generate dsRNAs from bi-directional promoters that are then processed by the RNAi machinery to promote heterochromatin formation (reviewed in (Martienssen and Moazed 2015)). DRIPc-seq data from *rnh1*Δ*rnh201*Δ cells indicated that RNAs originating from centromeric repeats were indeed immuno-precipitated by S9.6 (Figure 2C). These centromeric DRIPc-seq signals were resistant to exogenous RNase H treatment or *in vivo* expression of exogenous RnhA but sensitive to RNase III treatment, strongly suggesting that they corresponded to dsRNAs and not to RNA:DNA hybrids. We confirmed that RNase III did not affect R-loop formation and detection using *in vitro* substrates (Figure 2A). Taken together, these observations establish that S9.6 can immuno-precipitate *in vivo* produced and biologically relevant dsRNAs. Furthermore, these data strongly suggest that the affinity of the S9.6 antibody for dsRNAs can confound results of R-loop mapping approaches through DRIPc-seq type experiments, at least in fission yeast where significant loads of exosome-sensitive dsRNAs are produced.

### Improved R-loop mapping identifies two classes of R-loops

To map R-loops in the absence of contaminating dsRNAs, we treated genomic DNA with RNase III prior to S9.6 immuno-precipitation. Strikingly, DRIPc-seq signals after RNase III treatment exhibited all the hallmarks of genuine R-loops: they mapped to transcribed gene units, were strand-specific and co-directional with transcription, sensitive to the *in vivo* expression of exogenous RNase H (*E. coli* RnhA) and stabilized by loss of endogenous RNase H activity (*rnh1*Δ*rnh201*Δ) (Figure 3AB). Upon expression of RnhA, stranded DRIPc-seq signals were still detectable at ~25% of loci forming R-loops in wild-type cells, albeit with significantly reduced intensity and length (Figure 3BC). On the contrary, in the absence of endogenous RNase H activity (*rnh1*Δ*rnh201*Δ), both the number of R-loop forming loci and the percentage of the genome covered by R-loops increased significantly (Figure 3BC). Taken together, these data show that the mapping of R-loops by DRIPc-seq in fission yeast requires the elimination of dsRNAs by RNase III treatment. Unless further specified, all references to DRIPc from this point on refer to DRIPc-seq signal after RNase III treatment.

**Figure 3:**
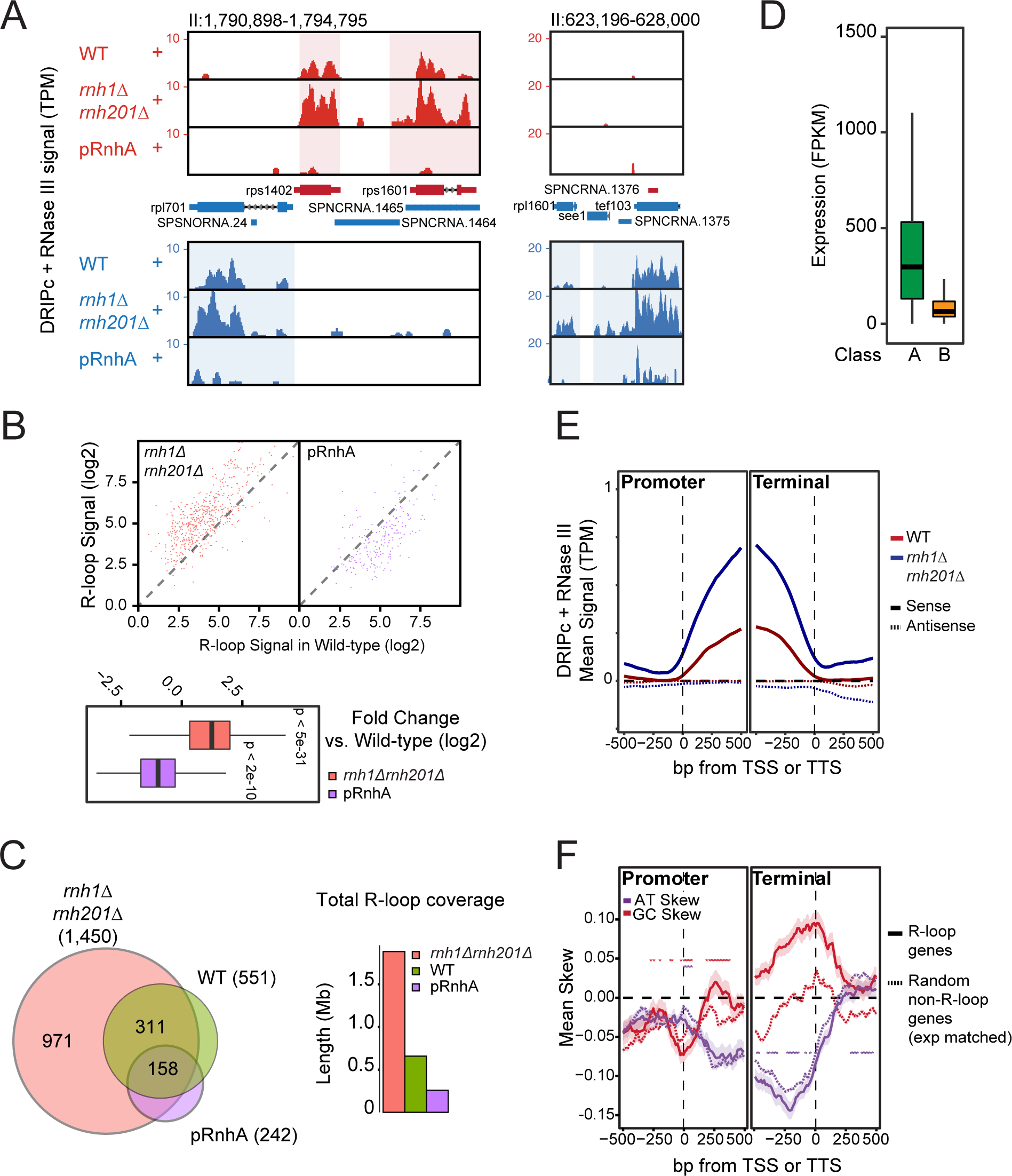
Improved R-loop mapping identifies two classes of R-loops in fission yeast. **(A)** Representative genomic regions showing DRIPc-seq signals after pre-treatment with RNase III. Color schemes are as in Figure 1A. Data is shown for WT, *rnh1*Δ*rnh201*Δ, and pRnhA cells. **(B)** Top: XY plot of RNase III-treated DRIPc-seq signal at each locus in WT (x-axis) vs. *rnh1*Δ*rnh201*Δ or RnhA expression (y-axis). This confirms that R-loop signals are stronger in *rnh1*Δ*rnh201*Δ and weaker in pRnhA for the vast majority of R-loop forming loci. Bottom: A box-plot representation of the same result. DRIPc-seq signals are significantly higher in *rnh1*Δ*rnh201*Δ and lower in pRnhA cells (p-values measured by Wilcoxon Mann Whitney test). **(C)** Left: Venn diagram showing the number of genes with R-loop peaks and their overlap in WT, *rnh1*Δ*rnh201*Δ and *pRnhA* conditions. Right: Barplot showing total DRIPc peak length in the indicated samples. **(D)** The wild-type transcript levels at genes with a class A or class B DRIPc-seq peak are represented as box-plots. Genes forming class B R-loops tend to be less expressed than genes forming class A R-loops. **(E)** Metaplots of RNase III-treated DRIPc-seq signals at promoter and terminator regions. **(F)** Metaplots of AT- and GC-skew at promoter and terminator regions of genes with class A R-loops and random non-R-loop forming genes matched for gene expression and length.

Our improved DRIPc-seq data identified two classes of R-loops, here referred to as class A and B, which differ mainly by their sensitivity to endogenous RNase H activity. Class A R-loops were reproducibly detected at ~500 loci in wild-type cells, representing roughly 5% of the 12,57 Mbp-long genome (Figure 3C). In cells lacking endogenous RNase H activity (*rnh1*Δ*rnh201*Δ), we detected an additional ~1,000 R-loop forming loci, which we refer to as class B R-loops. Overall, class B R-loops cover an additional ~10% of the genome. The average size of individual class A R-loops was longer than class B R-loops (Figure S4) and formed predominantly over highly transcribed genes and Tf2 retrotransposons, in a similar pattern to R-loop formation in *S. cerevisiae* (Chan et al. 2014; El Hage et al. 2014; Wahba et al. 2016) (Figure 3D and data not shown). Upon expression of bacterial RnhA, the few remaining detectable R-loops also formed at highly expressed genes, presumably from incomplete degradation of class A R-loops. On the contrary, class B R-loops formed at loci with weaker transcript levels (Figure 3D) and likely correspond to transcription units where transient R-loop formation is efficiently antagonized by endogenous RNase H enzymes. The heavily transcribed tDNAs, which accumulated class B R-loops in the absence of endogenous RNase H activity (Figure 2C and see (Legros et al. 2014)), were an exception to this low-expression rule for class B R-loops. The presence of R-loops at tDNAs confirms our previous observations (Legros et al. 2014) and observations made in budding yeast (Chan et al. 2014; El Hage et al. 2014; Wahba et al. 2016) and *Arabidopsis thaliana* (Xu et al. 2017). The low-level R-loop formation at tDNAs in wild-type cells is also consistent with the high recruitment of RNase H1 and Senataxin (Sen1) at these loci in fission yeast (Legros et al. 2014).

We did not detect a significant bias for R-loop formation towards either the promoter or the terminator regions of genes (Figure 3E) but we detected a significant strand bias for G over C (GC skew) in the terminal regions of class A R-loops (Figure 3F). By contrast, AT skew, which was identified as a feature of *S. cerevisiae* R-loops (Wahba et al. 2016), was not strongly different in R-loop-forming versus expression-matched non-R-loop genes (Figure 3F). Finally, we analysed whether R-loop forming genes associated with certain chromatin marks or factors compared to expression- and length-matched non-R-loop genes. R-loop-forming genes were significantly associated with a number of chromatin factors and marks compared to matched genes. The SMARCAD1 family ATP-dependent DNA helicase Fft3, the H3K4 methyltransferase Swd1, the H3K36 methyltransferase Set2, the histone chaperone Spt6 and the Paf1 complex subunit Ctr9^Tpr1^ were significantly enriched over R-loop forming genes (Figure S5). In conclusion, R-loop formation in fission yeast is widespread and associated with specific chromatin features characteristic of transcription elongation.

### RNase H level manipulations in fission yeast alters the transcriptome

Because we showed that manipulating RNase H levels *in vivo* affected R-loop formation, we assessed the impact of R-loop levels on the transcriptome of exponentially growing vegetative cells. For this, R-loops were either resolved by expression of bacterial RnhA in fission yeast cells, or stabilized by deletion of both endogenous RNase H genes, and the consequences on the transcriptome determined by strand-specific RNA-seq on ribo-depleted RNAs. RnhA expression impacted the steady-state levels of 1,218 transcripts out of 7,016, including 1,012 protein-coding mRNAs (Figure 4A). Of those protein-coding mRNAs, the abundance of 422 transcripts increased. Gene ontology (GO) analysis indicated that many up-regulated genes took part in ribosome biogenesis and nucleolus function (Figure 4A). Deletion of both endogenous RNase H genes was accompanied by significant changes in expression of 660 transcripts, including 584 protein-coding genes (Figure 4A). Many up-regulated genes again showed strong enrichment for GO terms related to ribosome assembly and nucleolus biogenesis. Further analysis revealed a significant overlap between the transcripts impacted by RnhA expression (loss of R-loops) and those affected by RNase H deletions (R-loop stabilization) (Figure 4B). For instance, genes involved in ribosome biogenesis and assembly tended to be up-regulated in both conditions, while genes involved in iron assimilation and glycan metabolism tended to be down-regulated. This unexpected observation suggests that long-term R-loop level manipulation through RNase H deletion or over-expression strategies impart significant and specific transcriptional responses in fission yeast.

**Figure 4.**
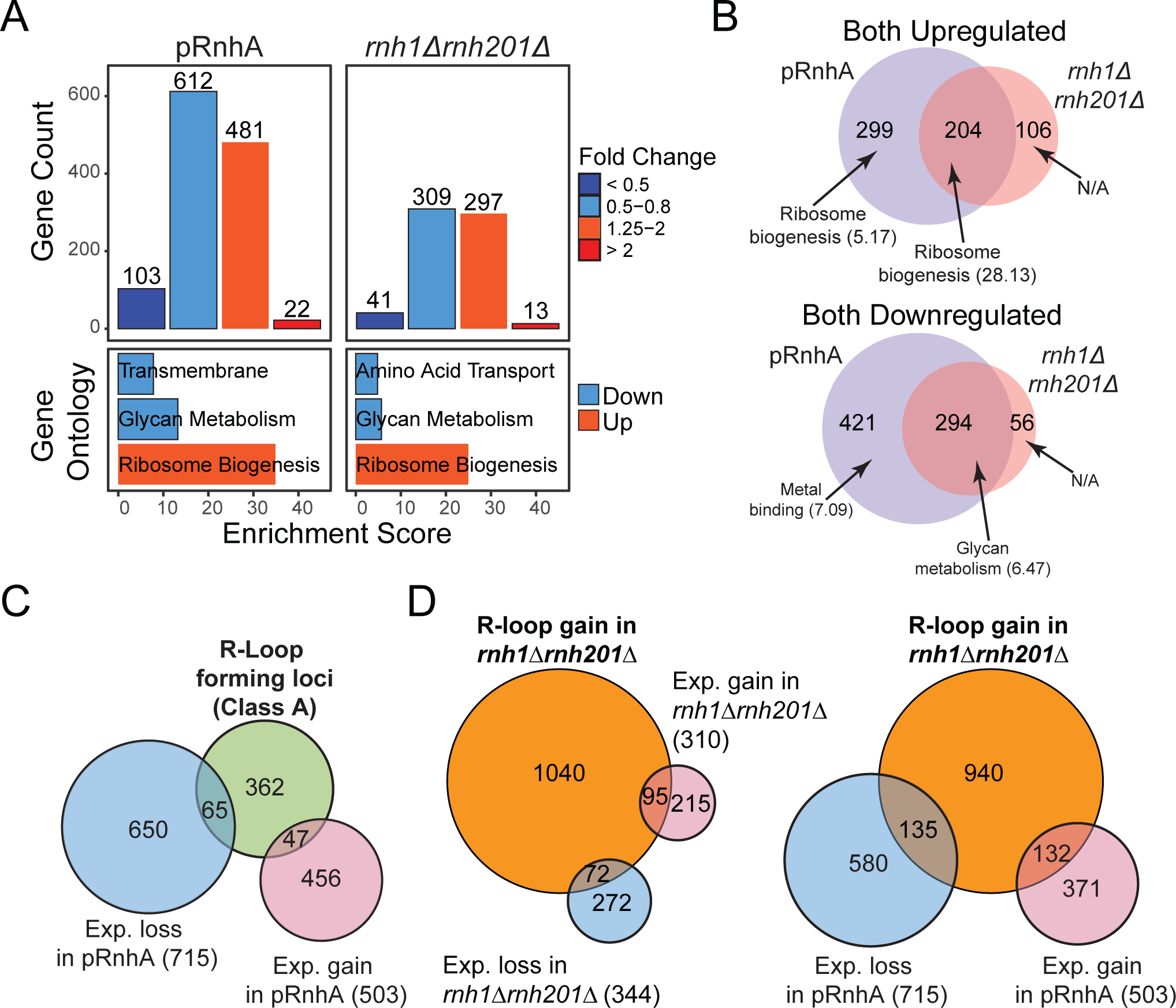
Manipulations of R-loop levels impact transcription predominantly at genes that do not form R-loops. **(A)** Top: Histogram showing the number of transcripts that were downregulated and upregulated after expression of RnhA (pRnhA) or deletion of both endogenous RNase H enzymes (*rnh1*Δ*rnh201*Δ). Bottom: Gene ontology analysis of up- and down-regulated transcripts. **(B)** Venn diagram showing the overlap between the transcripts that are up- (top) and down-regulated (bottom) in *pRnhA* and *rnh1*Δ*rnh201*Δ. GO term enrichment are indicated **(C)** Venn diagram showing the overlap between the transcripts forming class A R-loops and the transcripts whose levels are affected upon RnhA expression. **(E)** Venn diagram showing the overlap between the transcripts forming R-loops in *rnh1*Δ*rnh201*Δ and the transcripts whose levels are affected in *rnh1*Δ*rnh201*Δ (left) or upon RnhA expression (right).

### RNase H level manipulations in fission yeast alters predominantly the expression of genes that do not form detectable R-loops

To further understand the relationship between R-loop level perturbations and the associated transcriptional response, we looked for R-loop forming transcripts whose expression was impacted by RNase H over-expression or deletion. Surprisingly, the expression of less than a quarter of R-loop forming genes in wild-type cells (class A) was affected by RnhA expression (pRnhA, Figure 4C). Conversely, only about 10% of genes showing significant gene expression changes upon RnhA expression were R-loop forming in wild-type cells (Figure 4C). Similar results were obtained when focusing on class B R-loops (Figure 4D): transcript abundance at genes that gain R-loops upon RNase H deletion (class B) were for the most part not affected by loss of RNase H (Figure 4D, left panel) or gain of RNase H (Figure 4D, right panel). This shows that while R-loop level manipulation caused significant alterations to the transcriptome, the vast majority of responsive genes did not form R-loops that we could detect using DRIPc-seq. Taken together, our results suggest that the manipulation of RNase H levels in fission yeast triggers a global transcriptional response that impacts predominantly transcripts involved in ribosome biogenesis, irrespective of whether or not they can form detectable R-loops.

## DISCUSSION

### Caveats associated with the use of the S9.6 antibody to map R-loop forming regions

We provide evidence that the S9.6 antibody used in all variants of the DNA:RNA Immuno-precipitation (DRIP) method to map R-loops can also recognize and immuno-precipitate RNase III-sensitive double-stranded RNAs (dsRNAs) both *in vitro* and *in vivo* in fission yeast (Figures 1&2). As a result, it is likely that S9.6 can also recognize structured single-stranded RNA molecules that form hairpins. Our work shows that the affinity of S9.6 for dsRNAs becomes a problem when DRIP is followed by RNA sequencing for increased resolution, such as in DRIPc-seq, because it can lead to mixed signals from R-loops and dsRNAs. This is true in organisms or genotypes where significant dsRNA loads are produced. Recent DRIPc-seq maps in mammalian cells were strictly directional even in the absence of RNase III treatment (Sanz et al. 2016), suggesting that dsRNA formation was not an issue, at least in the conditions tested. Based on our *in vitro* results, it is also conceivable that even when DRIP is followed by direct DNA sequencing or qPCR, such as in classic DRIP-seq or DRIP-qPCR, competition from dsRNAs could affect the efficiency of the S9.6 immunoprecipitation (Figure 2). To circumvent this issue and ensure the quantitative enrichment of R-loops with the S9.6 antibody, we would recommend the use of RNase III treatments prior to immuno-precipitation.

DRIPc-seq data prior to RNase III treatment suggest that exosome-sensitive dsRNA species form widely in fission yeast, arguing for widespread antisense transcription. Such dsRNAs were not observed by regular RNA-seq experiments, suggesting that they most likely correspond to low-abundance RNA species that are detected after enrichment by the S9.6 antibody. dsRNAs were also recently detected in the distant budding yeast (Wery et al. 2016), suggesting that widespread dsRNA formation is a conserved phenomenon, at least in yeasts. dsRNA formation in fission yeast is associated with gene silencing through heterochromatin formation via the RNAi pathway (Martienssen and Moazed 2015). Our data indicate that the percentage of the genome to which these dsRNAs map is a lot greater than the percentage of the genome covered by heterochromatin. Further investigation will be required to establish whether or not such widespread low-level dsRNA formation plays a biological role.

### Improved R-loop mapping in fission yeast

The R-loop maps provided here represent a significant improvement over the two maps that were published previously in fission yeast. The first map was generated by DRIP-seq and as a result was not stranded and had low resolution (Castel et al. 2014). The second map was made using a ChIP-exo approach to increase resolution but it only provided a map of R-loops in cells lacking RNase H activity (*rnh1*Δ*rnh201*Δ cells) and the sensitivity of the signals to RNase H was not controlled (Ohle et al. 2016). The maps produced here are stranded, have high resolution, and the sensitivity of the signals to *in vivo* RNase H level manipulations has been validated.

While our maps recapitulate many observations made previously in yeast (R-loop formation at highly expressed G-rich genes, at Tf2 retrotransposons and at tDNAs), they differ from previous maps in several respects. For instance, contrary to what was proposed before (Castel et al. 2014; Ohle et al. 2016), we could not detect significant R-loop formation either over heterochromatin repeats at centromeres or over gene terminator regions in fission yeast. Although we confirmed that the S9.6 antibody immuno-precipitates RNA species associated with centromeric repeats and terminator regions (Figures 1&2 and Figure S2), our data suggest that these RNAs are resistant to both *in vivo* RNase H over-expression and *in vitro* RNase H treatment whilst being sensitive to RNase III treatment. As such, these RNAs are more likely to correspond to dsRNAs rather than R-loops. We speculate that at least a fraction of these dsRNAs remain associated with chromatin as was suggested previously for heterochromatin transcripts (Bronner et al. 2017), which might explain why they have been previously detected using DRIP. Interestingly, lack of a significant enrichment for R-loops at terminator regions was also recently observed in *Arabidopsis thaliana* (Xu et al. 2017). The situation in *S. pombe* and *A. thaliana* is therefore different from human and mouse where genuine R-loops form at a subset of terminator regions (Skourti-Stathaki et al. 2011; Ginno et al. 2012; Ginno et al. 2013; Skourti-Stathaki et al. 2014; Hatchi et al. 2015; Sanz et al. 2016).

Altogether, our improved R-loop maps show that 5% of the fission yeast genome can exist under an R-loop form in unperturbed cells, an observation consistent with findings in mammals and plants (Sanz et al. 2016; Xu et al. 2017). Under these conditions, R-loop formation is principally a property of highly transcribed genes (Class A). When cells lack endogenous RNase H activity, a second class of R-loops becomes detectable, covering an additional 8% of the genome. This shows that one of the roles of endogenous RNase H enzymes is to suppress low-level R-loop formation over a significant portion of the genome. Our data, combined with observations from mammalian cells and *A. thaliana* (Ginno et al. 2012; Ginno et al. 2013; Sanz et al. 2016; Xu et al. 2017) reinforce the idea that genuine R-loops share universal hallmarks, including strand-specificity, co-directionality with transcription, co-dependence with transcription levels, and a strong relationship with favourable DNA sequence features such as GC skew (Figure 3). In addition, we show that R-loop forming loci in fission yeast significantly associate with chromatin marks of active transcription and transcription elongation, consistent with properties of mammalian and plant R-loops (Sanz et al. 2016; Xu et al. 2017) These associations were significant even after stringent matching to non-R-loop forming genes with similar lengths and expression levels.

### Manipulation of R-loop levels over many cell generations impart significant transcriptome alterations

We report here that *in vivo* manipulations of R-loop levels trigger significant transcriptome alterations and that the responses to R-loop loss or stabilization are unexpectedly similar (Figure 4). This could be consistent with the idea proposed previously that the formation and the resolution of R-loops are equally important for gene expression (Skourti-Stathaki et al. 2011). How opposite changes in R-loop turnover rates are translated to similar gene expression responses remains unclear, however. Alternatively, we propose that the manipulation of R-loop levels over many generations might trigger a global cellular stress response. Our observation that up-regulated genes are enriched for functions associated with ribosome biogenesis and protein synthesis suggests that cells are coping with perturbations in the process of protein translation. An attractive, but as yet untested, hypothesis to account for this transcriptional response is that perturbations of R-loop levels negatively impact the transcription, processing, or export of R-loop prone rRNAs, ultimately interfering with ribosome assembly. Interestingly, perturbations of ribosome assembly were shown recently to result in the Spt6-dependent activation of ribosome-related genes in *S. cerevisiae* (Gomez-Herreros et al. 2017). If this feedback response also exists in *S. pombe,* it could explain why R-loop level manipulations result in the increased abundance of transcripts involved in ribosome assembly (Figure 4).

Regardless of its origin, this cellular response may mask the immediate impact of RNase H level manipulations on the expression of R-loop forming genes. Indeed, consistent with previous observations in *S. cerevisiae* (Chan et al. 2014), we observed that transcript levels of most R-loop forming genes were not significantly altered by RNase H level manipulations over long periods of time (Figure 4), A corollary to these observations is that collateral alterations to the transcriptome and the proteome could contribute to the phenotypes associated with prolonged RNase H1 over-expression, complicating the interpretation of such experiments. Although technically challenging, a suitable alternative to this approach could be to expose cells to high levels of RNase H1 for a very short period of time only.

To conclude, our work shows that the cross reactivity of the S9.6 for dsRNA and possible indirect perturbations to the transcriptome associated with prolonged manipulation of R-loop levels are two confounding parameters that must be taken into account in the design of experimental strategies to understand R-loop functions.

## METHODS

### Fission yeast strains

A complete list of all of the strains used in this study is given in Table S1. Standard genetic crosses were employed to construct all strains.

### R-loop mapping via DRIPc-seq

DRIP was performed as before (Legros et al. 2014) with the following modifications to the way the genomic DNA was extracted: 6.10^8^ cells were resuspended in 7.5 mL of pre-incubation buffer (20 mM citric acid, 20 mM Na_2_HPO_4_, 40 mM EDTA pH 8.0 to which 30 mM of ß-mercaptoethanol was added just before use) and incubated at 30°C for 10 min under vigorous agitation. Cells were then pelleted and resuspended in 3.5 mL of Sorb/Tris buffer (1 M Sorbitol, 50 mM Tris/HCl pH 7.4, 10 mM ß-mercaptoethanol containing 10 mg of Zymolyase 100T) and incubated at 30°C for 20 min under vigorous agitation then immediately transferred on ice. The resulting spheroplasts were washed in 15 mL of Sorb/Tris buffer and resuspended in 375 μL of NP buffer (1M Sorbitol, 50 mM NaCl, 10 mM Tris/HCl pH 7.4, 5 mM MgCl_2_, 1 mM CaCl_2_, 0.75% IGEPAL, 1 mM ß-mercaptoethanol) to which 3 μg/mL of RNase A and 700 μL of 10% SDS was added. After 1 hour at 37°C, phenol/chloroform and ethanol precipitation was used to purify the genomic DNA. The DNA was digested with restriction enzymes (EcoRI, SspI, BsrGI, XbaI, HindIII) as described before (Legros et al. 2014), in the presence of 2U of RNase III (Ambion) or 4U of RNase H (Roche). The digested DNA was then separated in two and each half was incubated overnight at 4°C with 40 μL of Protein A-coupled Dynabeads previously incubated with 10 μg of S9.6 antibody according to the manufacturer’s instructions. After extensive washes of the beads, the immuno-precipitated nucleic acids were pooled, purified as previously described (Legros et al. 2014) and heated for 1 min at 94°C to denature DNA:RNA hybrids. DNA molecules were then digested with DNase I for 1 hour at 37°C and the RNA molecules were recovered after ethanol precipitation. The RNA molecules were analysed and quantified using both Bioanalyzer (Agilent) and Quantus (Promega). Sequencing libraries were prepared using NEXTflex™ Rapid Directional mRNA-Seq Kit (dUTP-Based) (Bioo Scientific) from 10 ng of RNA, without Poly-A enrichment or RNA fragmentation. Paired-End sequencing (75 bp) was performed on a NextSeq500 (Illumina).

### Bioinformatics analysis

Both DRIPc-seq and RNA-seq reads were trimmed with fastq-mcf. DRIPc-seq reads were mapped to *Schizosaccaromyces pombe* genome version ASM294v2 (accession GCA_000002945.2) using Bowtie v2.1.0, and RNA-seq reads were mapped with Tophat v2.0.9. Gene sequence and annotation (version 4/2015) were downloaded from pombase. DRIPc-seq peaks were called using StochHMM as previously described in (Sanz et al. 2016). Reads were counted using htseq-count v0.6.0; RNA-seq reads were counted per annotated gene while DRIPc-seq reads were counted per DRIPc-seq peaks. For both RNA-seq and DRIPc-seq, significantly up and down-regulated genes and peaks were determined using DESeq2 v1.10.1, with threshold of at least 1.25x fold change compared to wild-type (higher or lower) and an adjusted P-value of 0.1 or less. Gene ontology analysis was performed using DAVID v6.8 with *S. pombe* as reference. All programs were run with default parameters unless mentioned. External ChIP-seq dataset were downloaded from GEO accession number GSE49575. To test if R-loop-forming genes are enriched or depleted over a specific factor, we performed shuffling test as described in (Sanz et al. 2016). Briefly, R-loop-forming genes were shuffled 125 times on a set of randomly chosen non R-loop-forming genes, matched for gene expression and length, and the results were used to calculate fold enrichment and empirical p-values.

### Dot blot analysis on genomic DNA

The genomic DNA was purified and digested with restriction enzymes as for the DRIPc (see above). After digest, the genomic DNA was quantified by qPCR using 3 independent primer pairs. DNA concentrations were then normalized between samples and a new round of qPCR with the same 3 independent primer pairs was used to confirm the correct normalization. 2 μL of DNA was spotted on a Hybond^TM^-N+ membrane (Amersham) and cross-linked twice with UV (0,12J). To quantify R-loop formation, the membrane was blocked in MES/BSA buffer (100 mM MES pH 6,6, NaCl 500 mM, 0,05% Triton, 2 mg/mL BSA) for 30’ hour at RT and 0,02 μg/mL of S9.6 antibody was added and incubated overnight. After extensive washes in MES/BSA buffer, exposure to a secondary antibody, ECL revelation was performed using SuperSignal West Femto Maximum Sensitivity Substrate (ThermoScientific) according to standard protocols. To quantify dsRNA or dsDNA, the membrane was first blocked in PBS 1X 0,02% Tween-20 containing 5% Milk and incubated overnight with the dsRNA-specific J2 antibody (SCICONS J2 tech, dilution 1/1000) or the dsDNA-specific (ab27156 Abcam, dilution 1/1000).

### In vitro *transcription* of *R-loops.*

750 ng of the pFC53 plasmid that contains the R-loop forming part of the *AIRN* transcription unit from mouse (Ginno et al. 2012) was transcribed for 30’ at 37°C in the sense orientation using the T3 RNA polymerase according to the manufacturer’s instructions. After inactivation of the polymerase at 65°C for 10’, the reaction was incubated for 2 hours at 37°C with 0,25 μg RNase A and 20 units of ApaLI to cleave the plasmid in two when needed (Figure 2B). When required 2 units of RNase H was added to the reaction. The reaction buffer was as follows: 50mM Potassium Acetate, 20mM Trisacetate, 10mM Magnesium Acetate, 100µg/ml BSA, pH 7.9. 10 μg of Proteinase K was then added for 30’ at 37°C before purifying the DNA using standard phenol/chloroform and ethanol precipitation techniques.

### Production of dsRNAs

4,5 μg of the pFC53 plasmid (see above) was transcribed for 30’ at 37°C in both orientations using both T3 and T7 RNA polymerases. After inactivation of the polymerase at 65°C for 10’, the reaction was incubated with 30 μg of RNase A for 20’ at 37°C. The plasmid DNA was then digested with DNase I (Thermo Fisher) according to the manufacturer’s instructions. dsRNAs were then purified using standard phenol/chloroform and ethanol precipitation techniques. RNase III was purchased from Ambion (reference AM2290) and digests were performed according to the manufacturer’s instructions.

### Immuno-precipitation of R-loops and dsRNA in vitro

The purified R-loops produced from 750 ng of pFC53 and the dsRNAs purified from 4,5 μg of pFC53 were incubated alone or in combination in 300 μL of interaction buffer (Tris 10 mM pH 8.0, NaCl 140 mM, 0,5% Triton). 100 μL were kept as Input (fraction IN, Figure 2B). The remaining 200 μL were incubated with 4 μg of S9.6 antibody coupled to 15 μL of Protein A-coupled Dynabeads at 4°C with agitation. After an hour incubation, the beads were collected and the unbound fraction (FT) was kept. The beads were washed 3 times with 1 mL of interaction buffer (5’ incubation each time) and then resuspended in 200 μL of incubation buffer. After 5’ incubation, the 200 μL were collected and kept as fraction W4. The beads were resuspended in 200 μL of elution buffer (50 mM Tris pH 8.0, 10 mM EDTA, 0,5% SDS) contaning 50 μg of Proteinase K and incubated at 55°C. After 30’, the beads were eliminated and the supernatant recovered (IP fraction). The IN, FT, W4 and IP fractions were then purified using standard phenol/chloroform and ethanol precipitation techniques and resuspended in 22 μL of Tris 10 mM pH 8,0. 6 μL were used for dot blot analysis with the S9.6 (DNA:RNA hybrids), J2 (RNA:RNA hybrids) or ab27156 (dsDNA) antibodies. 12 μL were run on a 0,8% agarose TBE 1X gel. The different fractions were then diluted 1/10,000 for qPCR analysis using the primers AIRN qL1 (5’-TCAGCACAACCAAGGATCAC-3’) and AIRN qR1 (5’-AGGGTGAAAAGCTGCACAAG-3’) to monitor the enrichment of the R-loop forming fragment and using the primers Amp 216 FW (5’-AACTTTATCCGCCTCCATCC-3’) and Amp 353 RV (5’- AAGCCATACCAAACGACGAG-3’) to monitor the enrichment of the control fragment.

### RNA-seq

RNAs were extracted as previously described (Vanoosthuyse et al. 2014) from biological triplicates (2.10^8^ cells per sample) and analysed and quantified using both Bioanalyzer (Agilent) and Quantus (Promega). Ribosomal RNAs were depleted from 2 μg of total RNAs. Strand-specific sequencing libraries were prepared from 2 ng of rRNA-depleted RNAs using the NEXTflex™ Rapid Directional mRNA-seq (Bioo Scientific) and sequenced on a HiSeq2500 Illumina (single reads 50 bp).

## DATA ACCESS

Our data have been deposited in GEO (reference **GSE101086**).

## ACKNOWLEDGEMENTS

The authors are extremely grateful to members of the Chédin lab for comments. This work was supported by the ‘‘Agence Nationale de la Recherche’’ [TRACC, CHX2011 to V.V.]; the “Comité du Rhône de la Ligue contre le Cancer 2016” to V.V.; the National Institutes of Health [GM120607 to F.C.]; and by a Howard Hughes Medical Institute International Student Research fellowship to S.R.H.

